# Three-dimensional visualization of moss rhizoid system by refraction-contrast X-ray micro-computed tomography

**DOI:** 10.1101/2022.07.07.499130

**Authors:** Ryohei Yamaura, Daisuke Tamaoki, Hiroyuki Kamachi, Daisuke Yamauchi, Yoshinobu Mineyuki, Kentaro Uesugi, Masato Hoshino, Tomomi Suzuki, Toru Shimazu, Haruo Kasahara, Motoshi Kamada, Yuko T. Hanba, Atsushi Kume, Tomomichi Fujita, Ichirou Karahara

## Abstract

Land plants have two types of shoot-supporting systems, root system and rhizoid system, in vascular plants and bryophytes. However, since the evolutionary origin of the systems are different, how much they exploit common systems or distinct systems to architect their structures are largely unknown. To understand the regulatory mechanism how bryophytes architect rhizoid system responding to an environmental factor, such as gravity, and compare it with the root system of vascular plants, we have developed the methodology to visualize and quantitatively analyze the rhizoid system of the moss, *Physcomitrium patens* in 3D. The rhizoids having the diameter of 21.3 μm on the average were visualized by refraction-contrast X-ray micro-CT using coherent X-ray optics available at synchrotron radiation facility SPring-8. Three types of shape (ring-shape, line, black circle) observed in tomographic slices of specimens embedded in paraffin were confirmed to be the rhizoids by optical and electron microscopy. Comprehensive automatic segmentation of the rhizoids which appeared in different three form types in tomograms was tested by a method using Canny edge detector or machine learning. Accuracy of output images was evaluated by comparing with the manually-segmented ground truth images using measures such as F1 score and IoU, revealing that the automatic segmentation using the machine learning was more effective than that using Canny edge detector. Thus, machine learning-based skeletonized 3D model revealed quite dense distribution of rhizoids, which was similar to root system architecture in vascular plants. We successfully visualized the moss rhizoid system in 3D for the first time.

## Introduction

Rooting structure of vascular plants is called the root systems consisting of individual roots. The root system provides a basis for supporting and anchoring the upper part of the plants for growth by uptaking water and minerals from substrate, such as soil. The root system adapts itself to surrounding soil environment changing its architecture [1]. Understanding how the plant root system adapts themselves flexibly to the surrounding environment is important for optimization of plant cultivation conditions under given environment.

An individual rhizoid of mosses is a multi-cellular filament [2], and its morphology is totally different from an individual root of vascular plants, which is an organ composed of multiple tissues. Nevertheless, the rhizoids of the Bryophyte, the first land plants, has functions of supporting and anchoring the upper part of the plants as well as uptaking water and minerals from soil, similarly to the roots of vascular plants. Therefore, whole rhizoids, which belong to one individual gametophore, with their architectural organization called “the rhizoid system” hereafter for convenience, which is commonly used in previous literatures as contrasted with the root system of the vascular plants [3].

One of the most important environmental factors that affect architecture of the rooting structures is gravity. And the rhizoids of mosses show positive gravitropism in darkness [4]. Considering these common functional characteristics, therefore, understanding the regulatory mechanism of organizing the rhizoid system and comparing it with that of the root system of vascular plants would give a clue to understand how land plants evolved this mechanism.

In order to understand adaptive mechanisms of the root and rhizoid systems to different environmental factors, it is necessary to visualize their morphology in 3D and to extract quantitative parameters describing its 3D morphology. Exploring the methodology of 3D morphological analysis of compact rhizoid system will provide a model system that facilitates to analyze that of the root system architecture of vascular plants as well. Hypergravity experiments have been performed using centrifuges to investigate effects of gravity on various physiological phenomena [5, 6], where it has already been reported that length of the rhizoids and a whole mass (dry weight) of the rhizoid system of a model moss, *Physcomitrium* (*Physcomitrella*)*patens* increased under hypergravity. Investigating responses of moss rhizoid system to altered gravitational conditions will lead to understand the mechanism of architectural regulation in mosses and in root system of vascular plants by gravity.

With the aim to understand adaptive mechanisms of the root system of vascular plants to environmental factors, X-ray micro-CT is developed. Industrial CT scanners or a scanning system employing microfocus X-ray source has been widely used to visualize roots of crop plants [7–13]. On the other hand, we have reported that refraction-contrast X-ray micro-CT using coherent X-ray optics enabled to visualize thin dried secondary roots of a model plant Arabidopsis having the thickness of its isosurface model at 20.9 μm on the average [14]. Such a higher spatial resolution imaging method can be applicable to visualize thin rhizoids of mosses.

Recently, we have performed “Space Moss” experiment on the International Space Station to understand effects of microgravity on the growth of *P. patens* [15]. In the present study, we have at first aimed to visualize its rhizoid system architecture in 3D by refraction-contrast X-ray micro-CT using specimens obtained by the Space Moss experiment. We also investigated the rhizoid system architecture under a hypergravity condition performed on the ground to examine effects of altered gravitational conditions. While we have manually traced the signal of the root in the obtained tomograms and made 3D models in the previous study [14], here we undertook automatic segmentation methods of the rhizoid area based on image processing techniques such as contour extraction as well as that based on machine learning. Images obtained by automatic segmentation was compared with the ground truth images obtained by manual segmentation and confusion matrices were formed. Accuracy of automatic segmentation was evaluated by comparing indicators such as F1 score and IoU, revealing that the automatic segmentation based on machine learning was more effective than that using contour extraction. Isosurface and skeletonized models of the rhizoid system were visualized in 3D for the first time. Three-dimensional visualization and digitalization of complex network of the rhizoid system will be also applicable to visualize such as 3D distribution of root hairs of vascular plants and hyphal architecture of mycorrhizal fungi interacting with roots, which will give insight into how architecture of rooting system of land plants is evolved under the Earth’s environment.

## Methods and materials

### Plant materials and growth conditions

Plant materials used in the present study was *Physcomitrium (Physcomitrella)patens* (Hedw.) Bruch et Schimp.. These were obtained by the Space Moss experiment which was performed from 2019 to 2020 (Run 1, 2, and 3) as well as by the hypergravity experiment performed on the ground. Growth conditions of the Space Moss experiment are as follows. Forty-eight gametophores were planted on the ground on an agar slab (W × D × H = 54 × 44 × 25 mm) containing BCD medium [16] in a polycarbonate growth chamber having the outer dimensions of W × D × H = 60 × 50 × 60 mm. Growth chambers were installed in the Plant Experimental Units (PEU), which was designed for experiments in the International Space Station [17]. Plants were kept refrigerated until the start of the experiment. PEUs were installed either in a microgravity compartment or on a centrifuge of to produce artificial gravity in the CBEF [18, 19] in the ISS. This artificial 1 × *g* condition is referred to as Space 1 × *g*. For the ground control experiments, PEUs were installed in the Cell Biology Experimental Facility (CBEF) at Tsukuba Space Center. Plants were illuminated from the top with LED matrix [19] with the light intensity of 30 μmol m^-2^ s^-1^ at the bottom center of the growth chamber. Plants were grown for 25-26 days at 25 °C. Run 1 and 2 experiments were performed from Jul. to Aug., 2019 and from Dec. 2020 to Jan., 2021, respectively. For the record, specimens of Space Moss experiment used in the present study were as follows: Ground 1 × *g*_SN17; Ground 1 × *g*_27, 82; Space 1 × *g*_005; Space 1 × *g*_003, B003, 004; Space μ × *g*_102.

Growth conditions of the hypergravity experiment were basically the same as described previously [5] using MK3 centrifuge (Matsukura Co., Toyama, Japan) with the ground control experiments. Plants were grown under 10 × *g* or 1 × *g* (as the control) for 25-26 days at 25 °C. For the record, specimens of hypergravity experiment used in the present study were as follows: Ground 1 × *g*_3; Ground 10 × *g*_1, 2.

### Chemical fixation and embedding of rhizoid system specimens

The moss plants cultivated on orbit (Run 1 and 2 experiments) were chemically fixed on orbit together with the agar medium in 100 mM sodium-phosphate buffer (pH 7.2) containing

2.5 % (v/v) glutaraldehyde (Polysciences, Inc., Warrington, USA), and were kept refrigerated until the start of embedding process on the ground (Fig. 1a). Specimens were embedded in paraffin primarily for X-ray micro-CT and optical microscopy while those were embedded in LR White resin primarily for electron microscopy.

**Fig. 1.**
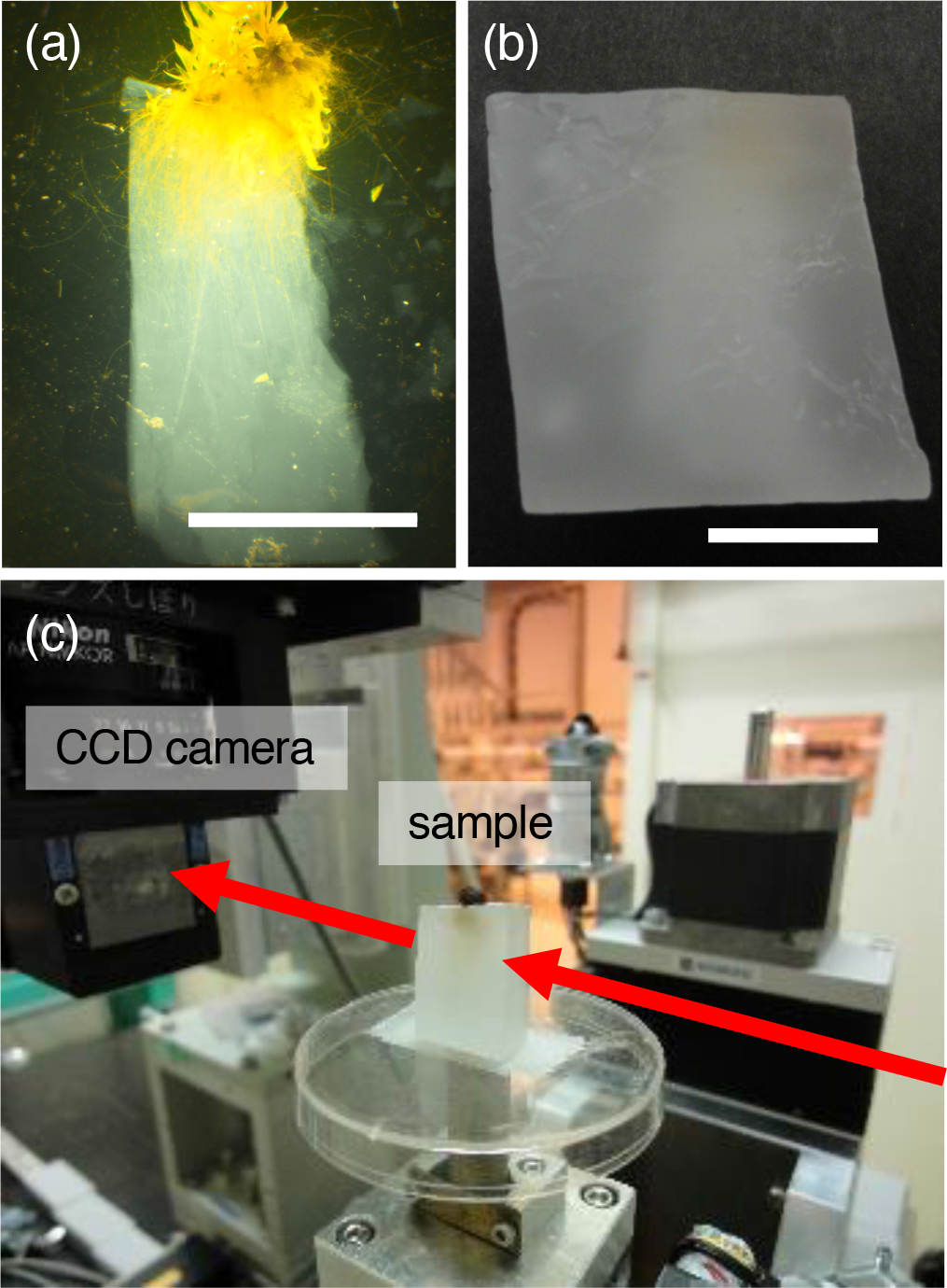
Specimen preparation and X-ray micro-CT scanning. (a, b) Lateral views of gametophores and rhizoids of *P. patens* grown on/in agar media observed under a dissecting microscope before (a) and after (b) embedding in paraffin. Specimen example: Ground 1 × *g*_SN17. Scale bars = 10 mm. (c) Experimental set up at the beamline BL20B2. Arrows indicate the X-ray path. Rhizoid system embedded in paraffin was placed on a rotation stage. A seed of morning glory was placed on the paraffin block as a position marker.

For specimen embedding in paraffin, fixed plant specimens of Run 1 were dehydrated in a graded ethanol series (70, 90, 99.9, 100 % (v/v)) and xylene in glass vials, which were placed on a rotary shaker (80 rpm). Specimens were then infiltrated with paraffin (Paraplast plus, Sigma-Aldrich Japan, Tokyo, Japan) for 3 days at 60 °C.

For specimen embedding in LR White resin (Electron Microscopy Sciences, Hatfield, USA), plant specimens of Run 2 chemically-fixed on orbit as mentioned above and plant specimens of hypergravity experiment (10 × *g* or 1 × *g* as the control) were fixed or refixed in 50 mM sodium-cacodylate buffer (pH 7.2) containing 2.0 % (v/v) glutaraldehyde and 1.0 % (w/v) paraformaldehyde (Wako Pure Chemicals, Osaka, Japan) at 4 °C for 12 h. The specimens were washed with DDW and were post-fixed at 4 °C for 3 h with 50 mM sodium-cacodylate buffer (pH 7.2) containing 1 % (w/v) osmium tetroxide. The fixed specimens were dehydrated in a graded ethanol series and were embedded in LR White resin. The fixed specimens were stained *en bloc* with EM stainer (Nisshin EM, Tokyo, Japan) during dehydration at the step of 10 % (v/v) ethanol.

### Optical microscopy of paraffin sections

Sections of 15 μm in thickness were cut using a rotary microtome from the specimens embedded in paraffin (Fig. 1b) and remained specimens were observed by X-ray micro-CT. Paraffin sections were mounted on slide glasses with gelatin adhesive, were extended at 40 °C on a hotplate, and were deparaffinized with graded xylene and ethanol series. Deparaffinized sections were stained with aqueous solution of 0.05 % (w/v) Toluidine blue O. Digital photographs were taken using a digital camera Cool Snap cf (Nippon Roper KK, Tokyo, Japan) or DS-Ri1 (Nikon, Tokyo, Japan) fitted to an optical microscope (BX-50, Olympus Corp., Tokyo, Japan).

### Electron microscopy of ultrathin sections

Ultrathin sections of 80 nm in thickness were cut using an ultramicrotome (EM UC7, Leica Microsystems GmbH, Wetzlar, Germany) from the specimens embedded in LR White resin, were stained with lead citrate for 1.5 min, and were observed under a transmission electron microscope (H-7650, Hitachi High-Tech Corp., Tokyo, Japan) at 80 kV.

### Refraction contrast X-ray micro-CT and tomographic reconstruction

Refraction contrast X-ray micro-CT was performed at the experimental Hutch 1 of the beamline BL20B2 of the SPring-8 synchrotron radiation facility at Japan Synchrotron Radiation Research Institute, according basically to the method described previously [14, 20]. Its experimental setup is shown in Fig. 1c and a brief overview of the method is as follows. The X-ray energy was adjusted to 15 keV. The images consecutively projected on the fluorescent screen were recorded by a CMOS camera (ORCA-Flash 4.0, Hamamatsu Photonics KK, Hamamatsu, Japan). Size of the obtained images was 2048 × 2048 pixels (ca. 5 × 5 mm). A series of 900 projections were recorded over 180 degrees. Because the rhizoids distributed deeper than 5 mm from the base of the gametophore, observation was separately done for the areas from the base to 5 mm below and from 5 mm to 10 mm below. The spatial (pixel) resolution of the 3-D structure was estimated to be 2.71 or 2.75 μm pixel^-1^. The convolution back projection method was used for tomographic reconstruction. Chesler filter, which is provided by the software package SP-μCT (http://www-bl20.spring8.or.jp/xct/), was used for reconstruction as previously determined [14]. Tomographic slices were obtained and 3-D models (isosurface, wireframe) were drawn using the IMOD software package [21] as previously described [22].

For comparing tomographic images with electron microscopic images, tomographic images had to be obtained particularly at the edge surface of a specimen just beneath where ultrathin sections were cut. However, structures near the edge surface of the specimen were hardly visible due to an artifact. To avoid this artifact, the edge surface of the specimen was covered with paraffin before scanning.

### Automatic segmentation of rhizoid area

Two methods were applied for automatic segmentation of cross-sectional areas of rhizoids (Fig. 2). One is based on contour extraction using Canny’s edge detection algorithm [23]. Contrast of the volume was normalized and equalized using Enhance Contrast function of Fiji software package (https://imagej.net/software/fiji/) [24] with saturated pixels at 0.3 %. Canny Edge Detector plugin of Fiji [25] was applied for each tomographic slice and contours were extracted from where intensity gradient is high with the following settings: Gaussian kernel radius at 2, Low threshold at 2.5, High threshold at 6. Then Closing (3D) plugin which connects interrupted rhizoid contours with the setting of Element Shape as Cube (X, Y, and Z radii are 1, 1, and 1, respectively) and Fill holes (3D) plugin which remove holes inside particles were applied. Both of them are MorphoLibJ (Release 1.5.0) plugins of Fiji [26].

**Fig. 2.**
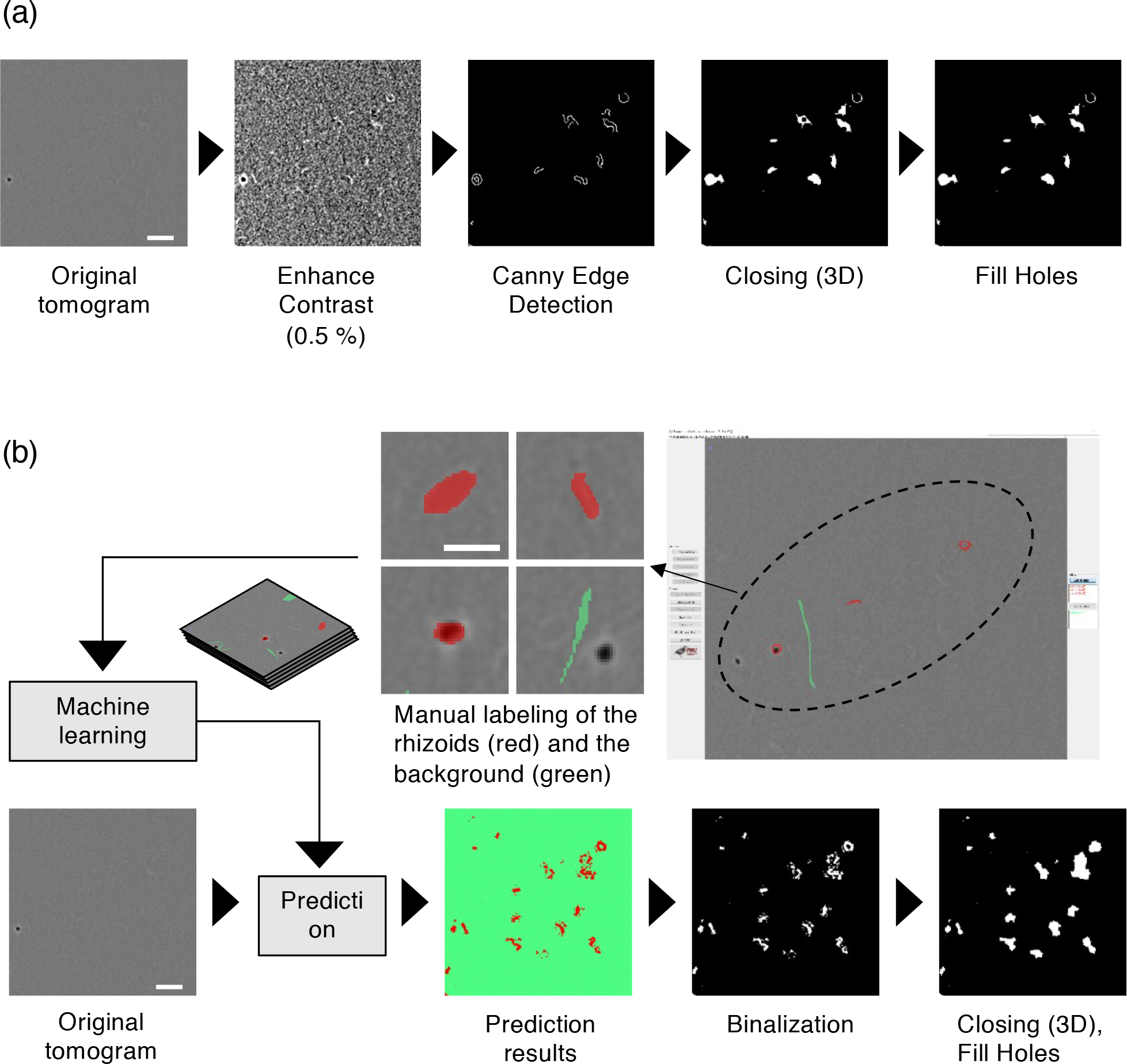
Workflow for automatic segmentation of cross-sectional areas of rhizoids. (a) A method based on contour extraction using Canny edge detector. (b) A method based on machine learning using TWS 3D. Specimen example: Ground 1 × *g*_82.

The other uses Trainable Weka Segmentation 3D (TWS 3D) plugin of Fiji software, which is based on pixel-based segmentation algorithms using machine learning [27]. In this program, pixels annotated manually by users are learned and a classifier is built by the machine learning platform WEKA (Waikato Environment for Knowledge Analysis) [28]. Mean and Variance filters, which extract features of the input images, and Fast Random Forest classifier were used as the default settings. A reconstructed volume was divided into substacks, each of which is composed of 100 transverse tomograms. In a representative substack, rhizoids and background pixels were manually annotated as independent classes in a part of a tomogram using Freehand selections tool and saved as a classifier model to train the classifier. This training was repeated for several times and the trained classifier was used to obtain a prediction result (Fig. 2b).

### Evaluation of the automatic segmentation methods

Automatically-segmented rhizoid areas by the two methods mentioned above were compared with manually-segmented areas to evaluate the results as follows. As examples, 9 different areas of 256 × 256 pixels sampled from 4 tomograms of the specimens of Ground 1 × *g*_27, Ground 1 × *g*_82, Space 1 × *g*_004, and Space μ × *g*_102. Rhizoid areas obtained by automatic segmentation were compared with those done by manual segmentation (ground truth). Pixels in an image were categorized into either of the following four groups, true positive (TP), false positive (fp), false negative (fn), or true negative (tn) (Supplementary Fig. 2), where a pixel in the rhizoid area is positive and that in the background area is negative in the ground truth image.

Calculations for the evaluation were performed using a program written in Python as follows. Automatically and manually-segmented images were transformed to matrices, where each pixel of the rhizoid and background areas in the images was assigned to values ‘1’ or ‘0’, respectively. In the resulting matrix which was obtained by a subtraction of an automatically-segmented image matrix from its corresponding manually-segmented image matrix (MS), an element as value ‘1’ or ‘-1’ was judged as fn or fp, respectively. Matrices showing distribution of fn and fp were designated as FN and FP, respectively. A matrix showing distribution of tp, TP, was obtained by a subtraction of FN from MS. A matrix showing distribution of tn, TN, was obtained by subtracting the FN, FP, TP from MS. A confusion matrix was constructed by counting the number of elements in each matrix. Accuracy, F1 score which indicates how accurately rhizoid areas are labeled, and Intersection over Union (IoU) which indicates how similar predicted rhizoid area is to the ground truth were calculated using formulas shown in Supplementary Fig. 2e.

### Skeletonization of automatically-segmented rhizoid area

An isosurface model was made from a part of an image stack file obtained by automatic segmentation of rhizoid areas using ‘imod auto’ command in the IMOD software package. Skeletonization of the isosurface models of the rhizoids was done using Skeletonize (2D/3D) plugins of Fiji software.

## Results

### Observation of specimens embedded in paraffin by X-ray micro-CT

On the overview of a cross-sectional tomogram many structures can be seen (Fig. 3a). These structures are considered to be rhizoids, which needed to be verified. When a closer look was taken at a cross sectional tomogram of the rhizoid system embedded in paraffin, three types of filamentous structures were observed (Fig. 3b-d). The first one, which is referred to as Type A structure, appeared ring-shaped (Fig. 3b). The second one, which is referred to as Type B structure, appeared linear (Fig. 3c). The 3rd one, which is referred to as Type C structure, is a void space appeared as a black circle with a halo (Fig. 3d). However, in some cases, even one filamentous structure appeared in different forms from Type A thorough Type C depending on the location when viewed in 3D (Supplementary Fig. 1). From this observation, we inferred that these types of filamentous structures are different forms of rhizoids.

**Fig. 3.**
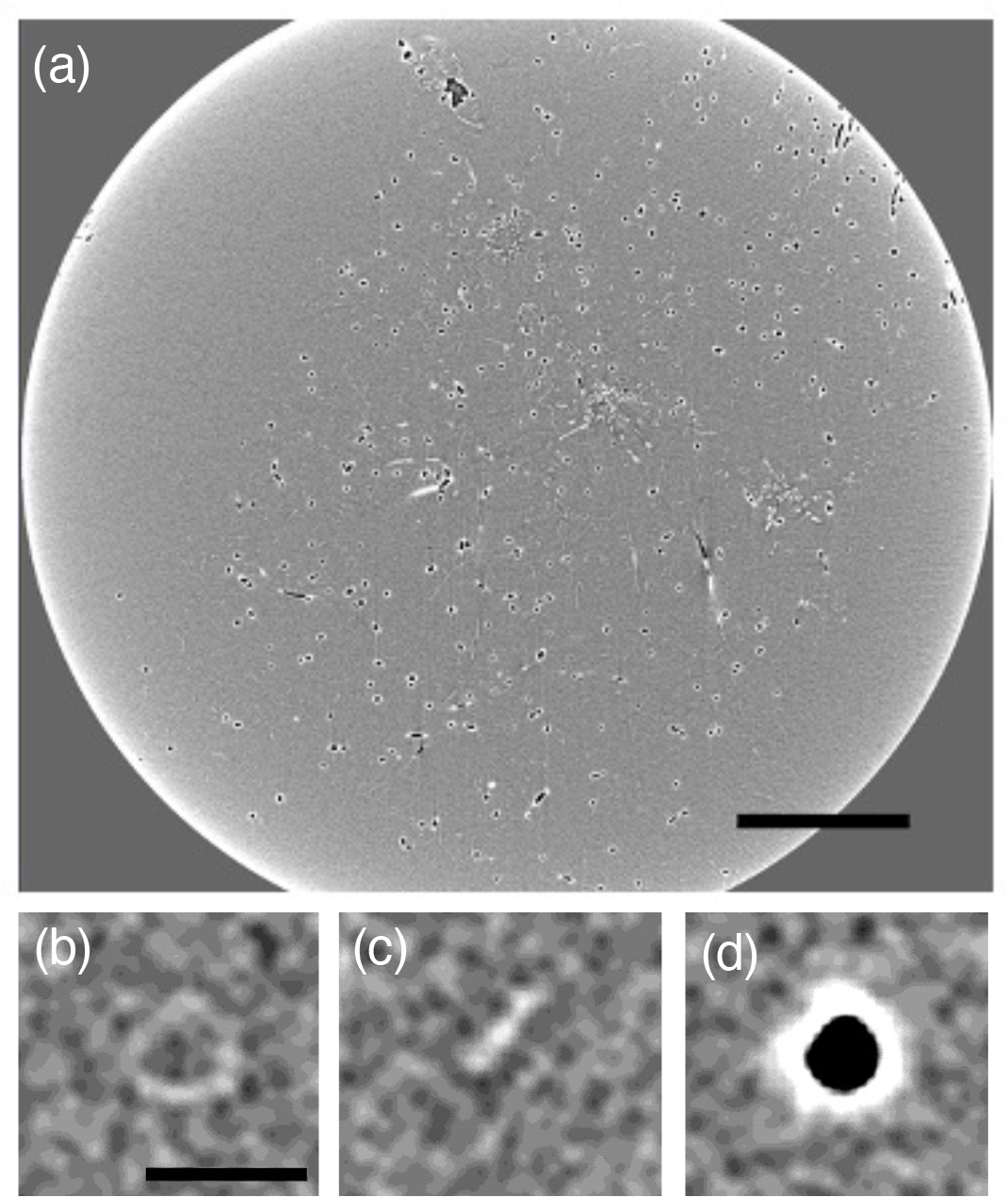
Observation of the reconstructed volume of the rhizoid system of *P. patens* embedded in paraffin. (a) Overview of a cross-sectional tomogram. (b-d) Cross sections of the three types of filamentous structures observed in the tomographic slices. (b) Type A, (c) Type B, and (d) Type C structures. Specimen example: a, Space 1 × *g*_003; b-d, Ground 1 × g_27 Scale bars = 1 mm (a), 50 μm (b-d).

### Comparison of tomograms and optical microscopic images of the three types of the filamentous structure

To test this hypothesis and to understand why rhizoids take such different forms, tomographic images and their corresponding optical microscopic images of the three types of the filamentous structures were compared. For this test, paraffin-embedded specimens of the hypergravity experiment performed on the ground were used instead to save the entire forms of irreplaceable specimens of Space Moss experiment. After reconstructing volumes by X-ray micro-CT, the three types of filamentous structures were observed on cross-sectional tomograms. Then actual cross-sections were cut from the paraffin-embedded specimens using a microtome at the corresponding positions of the tomograms and were observed under an optical microscope.

As a result, Type A structure, which appeared circular in tomograms, also appeared circular in the corresponding optical microscopic images (Fig. 4a). And Type B structure, which appeared linear in tomograms, appeared flat in the corresponding optical images (Fig. 4b). On the other hand, interestingly, Type C, which appeared as void spaces in tomograms, actually appeared flat similarly to Type B in the corresponding microscopic images (Fig. 4c). This means that Type C is actually the same as Type B. These observations raised questions why the filamentous structure take the form of circular or flat and why the void space, which appeared as Type C, was formed.

**Fig. 4.**
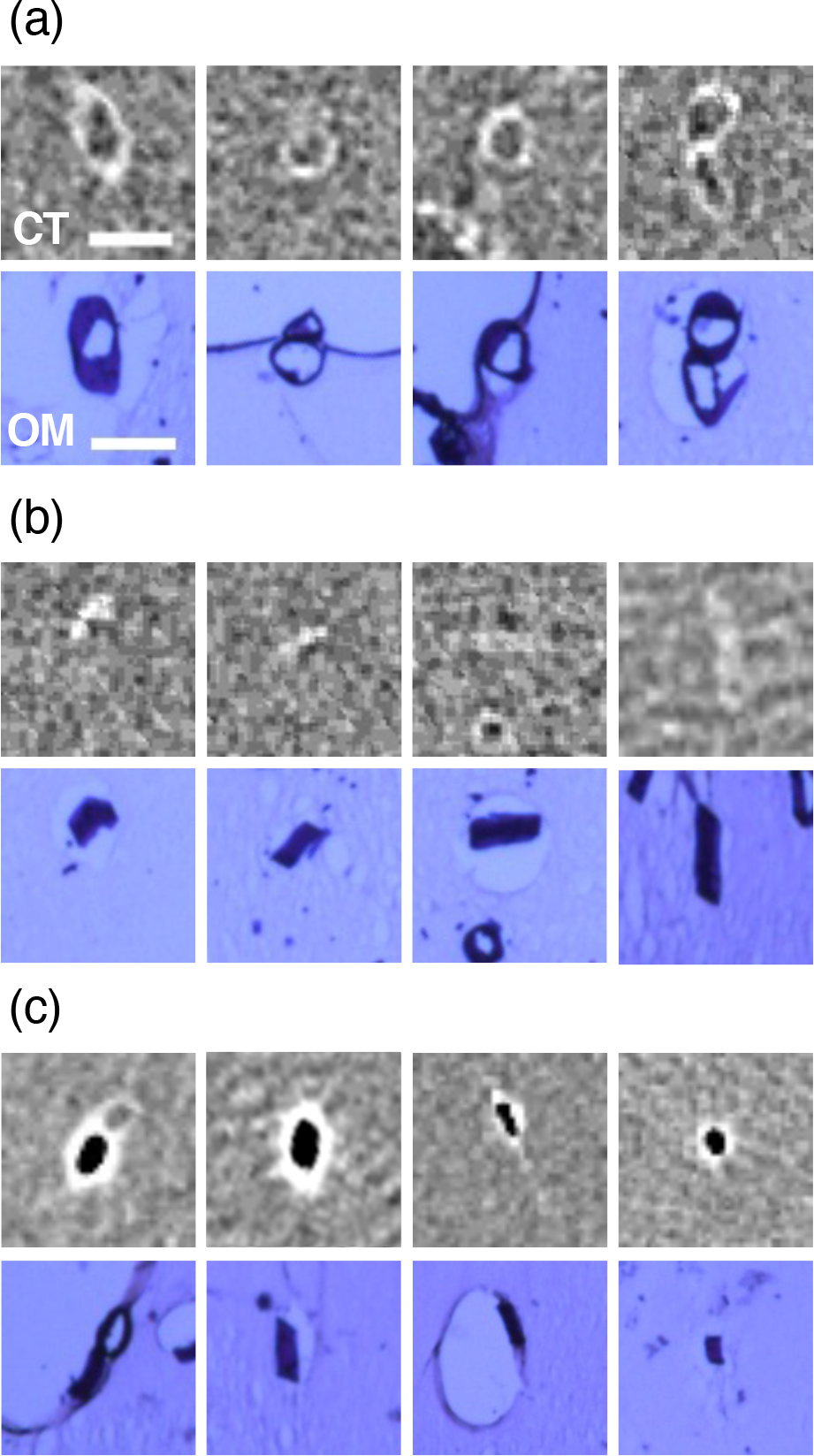
Comparison of tomographic images and optical microscopic images of the three types of the filamentous structures observed in the rhizoid system of *P. patens* embedded in paraffin. (a) Type A, (b) Type B, and (c) Type C structures. The upper row, CT: tomographic images. The lower row, OM: optical microscopic images of paraffin sections obtained at the position corresponding to its tomographic images. Specimen example: Ground 10 × *g*_1, 2. Scale bars = 50 μm.

### Observation of specimens embedded in LR White resin by X-ray micro-CT and under electron microscope

To answer the questions mentioned above, X-ray micro-CT was performed to reconstruct volumes using specimens embedded in LR White resin and tomograms were observed. As a result, however, only Type A was found while Type B and C were not (Fig. 5a). From this observation, we concluded that Type B and C are artifacts formed during paraffin embedding, which is probably due to a problem of infiltration of paraffin into the cell. Therefore, possible causes of formation of Type B and C are suggested as follows: a cell without infiltrated paraffin collapsed before paraffin solidification in the case of Type B, or collapsed after volatilization of xylene from the cell in the case of Type C. Although the flat structure is considered to remain adjacent to the void space in the case of Type C, an artifactual strong halo generated due to tomographic reconstruction around a void space in a tomogram might have made the structure hardly visible.

**Fig. 5.**
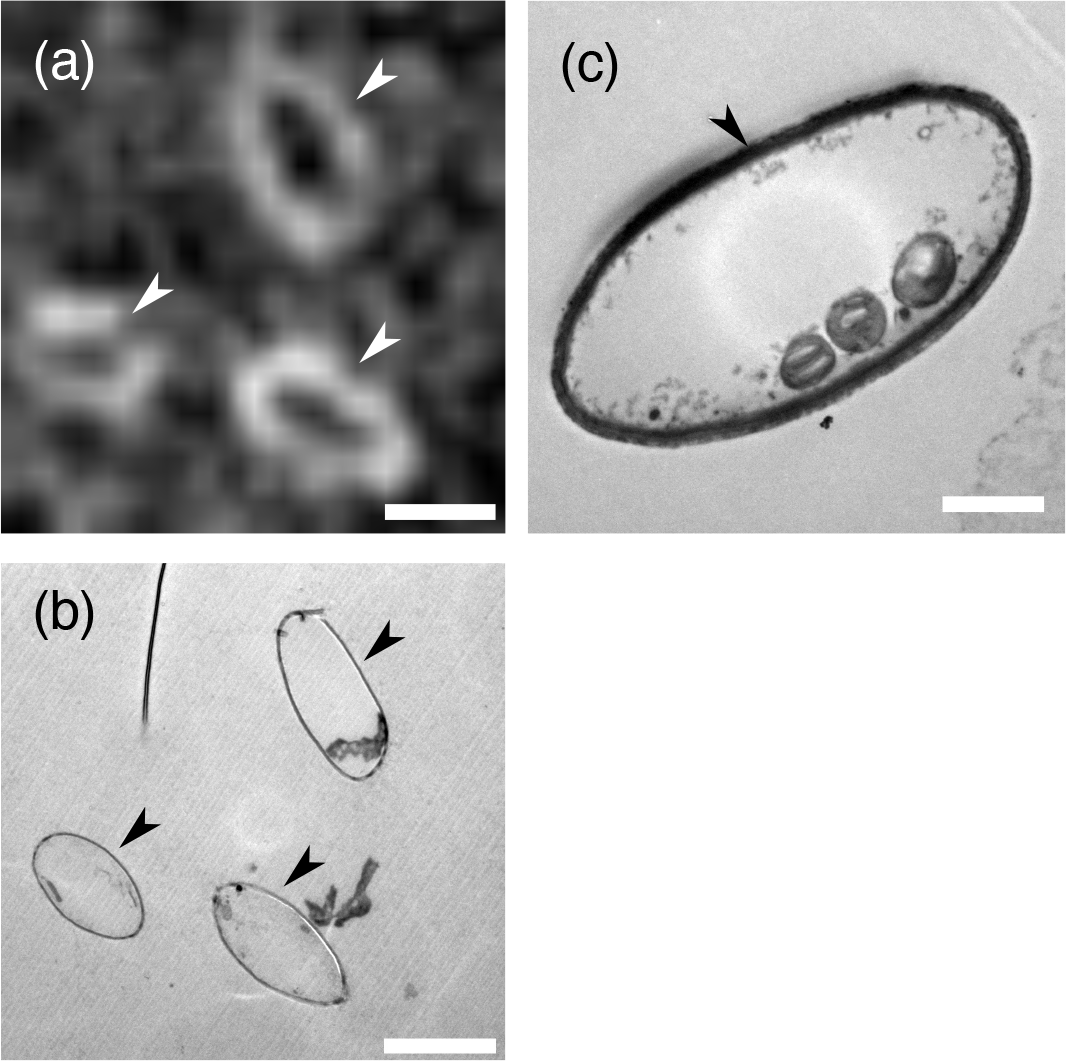
Comparison of a tomographic image and electron microscopic images of Type A structure observed in the rhizoid system of *P. patens* embedded in LR White resin. (a) A tomographic image, where three Type A structures are visible (arrowheads). (b) An electron microscopic image obtained at the position corresponding to the tomographic image of (a), where the same three structures are visible (arrowheads). (c) A magnified electron microscopic image of Type A structure, i.e., cross-sectional view of a rhizoid cell. An arrowhead shows cell wall. Specimen examples: Space 1 × *g*_B003 (a, b), Space 1 × *g*_004 (c). Scale bars = 20 μm (a, b), 5 μm (c).

Ultrathin cross-sections were cut from specimens embedded in LR White resin using the ultramicrotome from the same specimens at the corresponding positions of the tomograms. The ultrathin sections were observed under the transmission electron microscope (Fig. 5b, c). A magnified ultrastructural view clearly shows a cellular structure with organelles (plastids) surrounded by the cell wall (Fig. 5c), confirming that the filamentous structure is the rhizoid. White contours and darkly-stained contours of the circular structure (Type A) (Fig. 4a) were revealed to be the cell walls of the rhizoid (Fig. 5c, arrowhead).

Quantitative measurements showed that the rhizoids having the diameter of 21.3 ± 0.6 (μm, mean ± SE, n = 9) or the thickness of 5.2 ± 0.3 (μm, mean ± SE, n = 8) in the case of circular ones or flattened ones, respectively, were visualized by X-ray micro-CT, which were quantified using tomograms of Ground 10 × *g*_1 specimen.

### Segmentation and 3D modeling of the rhizoids

Manual segmentation of cross-sectional areas of rhizoids was tried at first to see distribution of rhizoids in a tomographic slice (Fig. 6). Because rhizoids were abundantly observed, it was unrealistic to perform segmentation of all rhizoids manually for every slice through the reconstructed volume. Therefore, we tested two methods of automatic segmentation of cross-sectional areas of rhizoids to label the three types of the rhizoid all together: one is based on contour extraction using Canny edge detector (Fig. 2a) and the other is based on machine learning using TWS 3D (Fig. 2b).

**Fig. 6.**
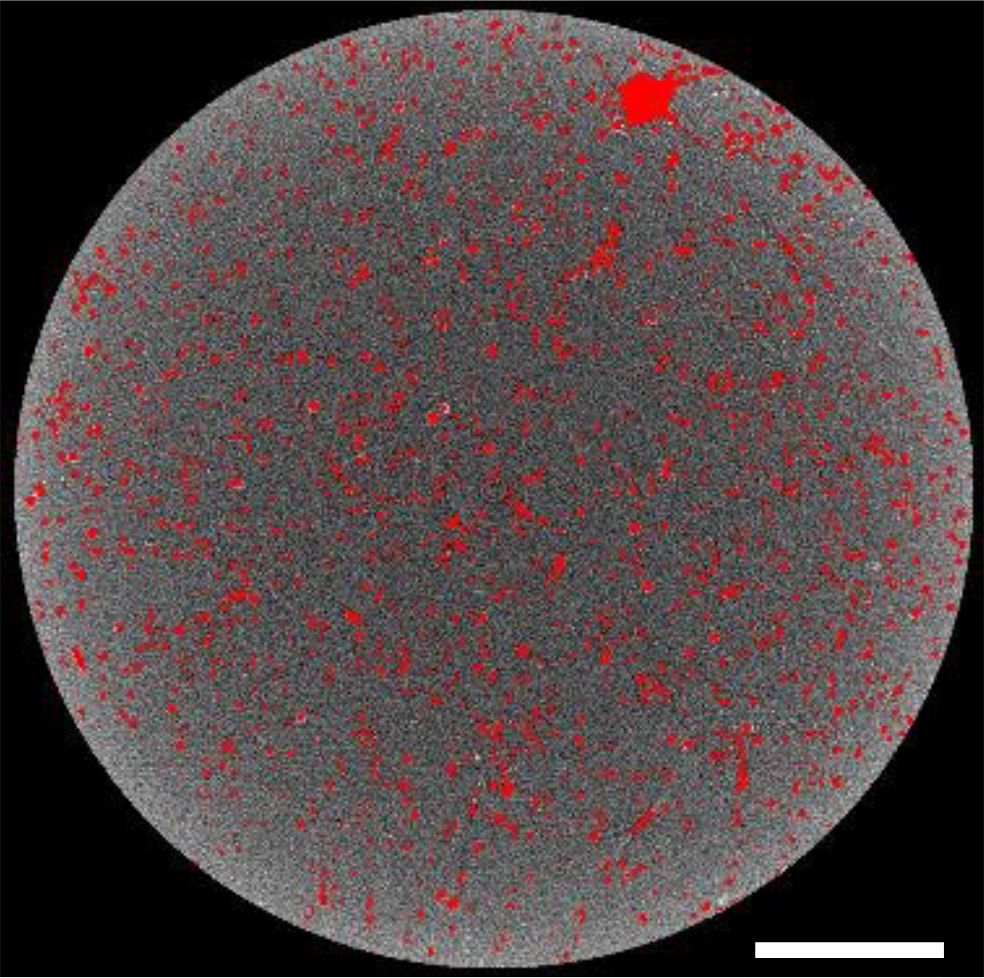
An overview of the rhizoid distribution in a cross-sectional tomographic slice of the rhizoid system of *P. patens* embedded in paraffin. Cross sectional areas of rhizoids are manually labelled in red. Specimen example: Ground 1 × *g*_27. Scale bar = 1 mm.

As a result, manually-segmented cross-sectional areas of rhizoids appeared in the form of the three types (Type A, B, and C) in tomograms (Fig. 7a, b). Rhizoids are also segmented by either automatic method successfully (Fig. 7c, d). However, cross-sectional areas of rhizoids segmented automatically using TWS 3D (Fig. 7d) appeared slightly larger than the manually-segmented ones (i.e., the ground truth) (Fig. 7b).

**Fig. 7.**
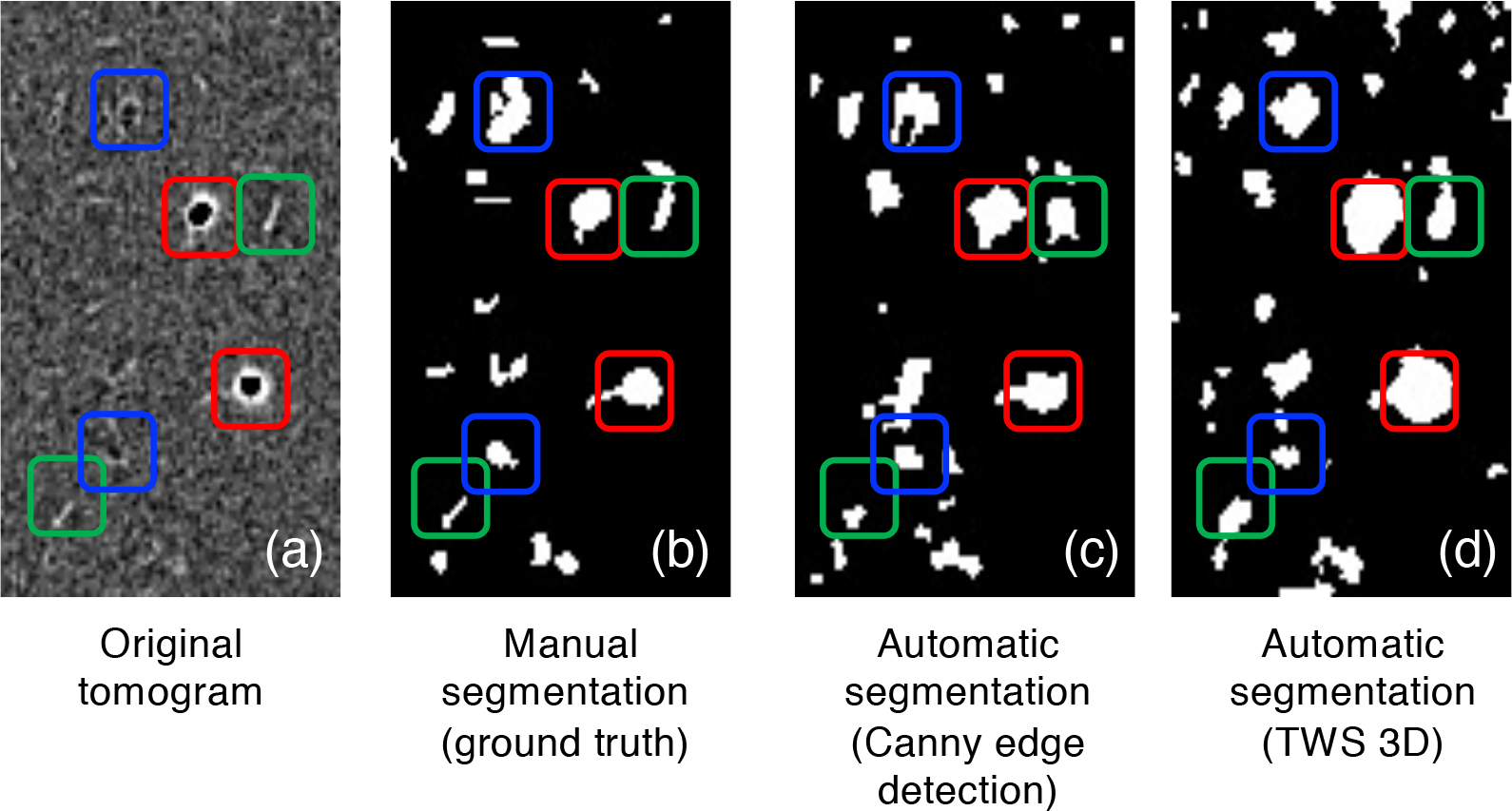
Comparison of manual and automatic segmentation of cross-sectional areas of rhizoids that appeared in the form of the three types in tomograms. (a) Original tomogram. (b) Manual segmentation (ground truth). (c) Automatic segmentation based on contour extraction using Canny edge detector. (d) Automatic segmentation based on machine learning using TWS 3D. Rounded squares indicate Type A (blue), Type B (green), and Type C (red). Specimen example: Ground 1 × *g*_27.

Automatically-segmented rhizoid areas by the two methods, including all of the three types (Type A, B, and C), were compared with manually-segmented ones. Pixels in automatically-segmented rhizoid areas were categorized into either of the following four groups, true positive, false positive, false negative, or true negative, and a confusion matrix was constructed by summarizing number of pixels in tomograms (Table 1). The confusion matrix actually showed that the number of false positive pixels is about three times larger in the result of the TWS 3D method than that of the Canny edge detector method while the number of false negative pixels in the result of the former is about half as large as that in the result of the latter.

**Table 1.**
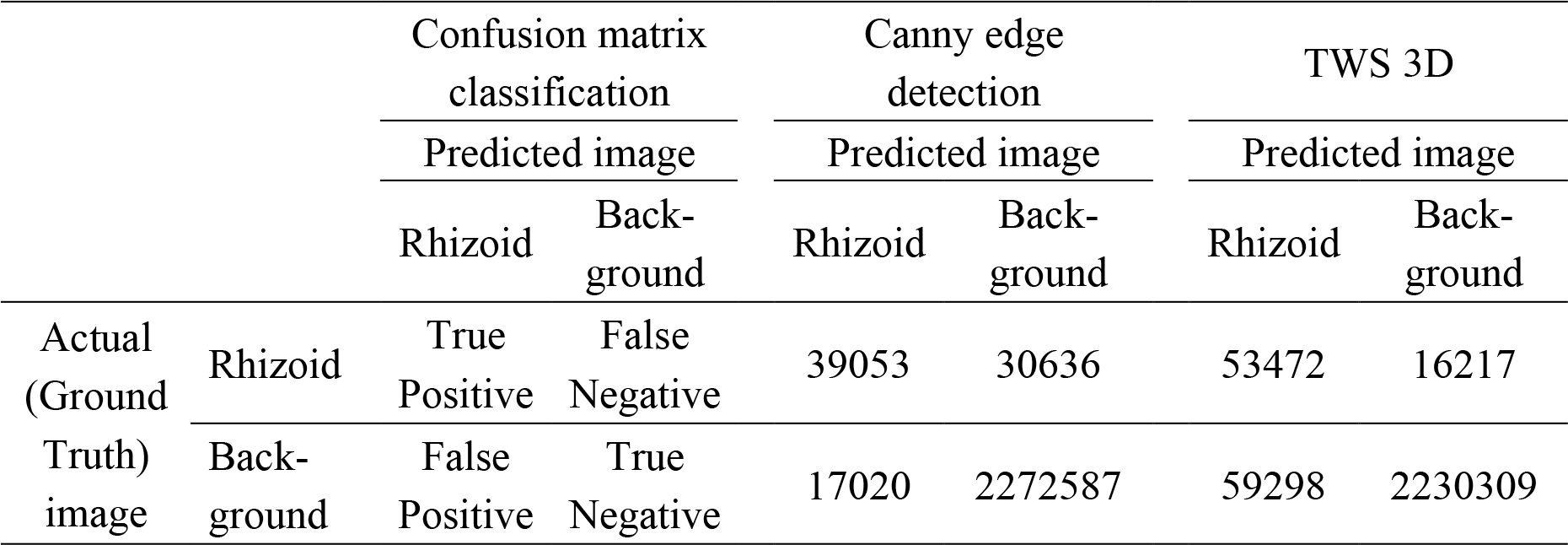
Confusion matrix results for evaluating two automatic segmentation methods. Numbers are summary of pixel numbers counted on 9 different areas of 256 × 256 pixels sampled from 4 tomograms of the specimens of Ground 1 × *g*_27, 82, Space 1 × *g*_004, and Space μ × *g*_102.

|  |  | Confusion matrix classification |  | Canny edge detection |  | TWS 3D |  |
| --- | --- | --- | --- | --- | --- | --- | --- |
|  |  | Predicted image |  | Predicted image |  | Predicted image |  |
|  |  | Rhizoid | Back-ground | Rhizoid | Back-ground | Rhizoid | Back-ground |
| Actual (Ground Truth) image | Rhizoid | True Positive | False Negative | 39053 | 30636 | 53472 | 16217 |
|  | Back-ground | False Positive | True Negative | 17020 | 2272587 | 59298 | 2230309 |

The two automatic segmentation methods were evaluated by calculation of accuracy, F1 score, and IoU (Fig. 8). The rhizoid area included all of the three types (Type A, B, and C). As a result, although no significant difference was found between the two automatic segmentation methods for accuracy, measures for performance evaluation such as F1 score and IoU are significantly higher in the result of the TWS 3D method than the Canny edge detection method. Because the area covered by the rhizoids are much smaller than the background, F1 score and IoU are important in the present case. Therefore, the method using TWS 3D was adopted for further analysis.

**Fig. 8.**
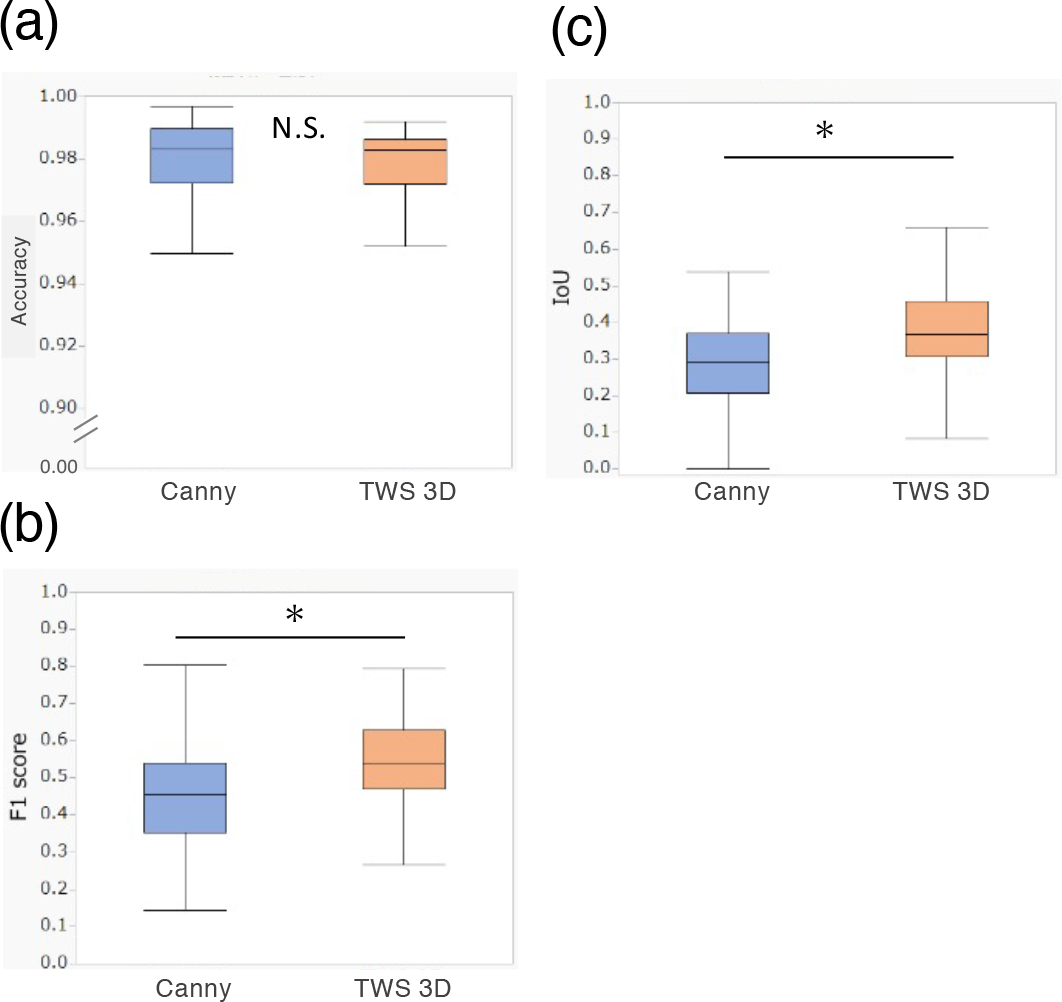
Comparison of accuracy (a), F1 score (b), and IoU (c) values between the automatic segmentation method based on contour extraction using Canny edge detector and the one based on machine learning using TWS 3D. N. S., Not significant (*P* ≥ 0.05); *, *P* < 0.05 (Welch’s *t*-test). n = 36.

### Three-dimensional architecture of the rhizoid system

Three-dimensional isosurface models of the rhizoids were made from automatically-segmented rhizoid areas obtained by application of the TWS 3D method (Fig. 9a, b). Distribution density of rhizoids appears to decrease with distance from the base of the rhizoid system (Fig. 9a). To see the distribution of the rhizoids clearly in 3D, the isosurface models were skeletonized (Fig. 9e). A lower magnification view shows a quite dense distribution of the rhizoids in 3D near the base (Fig. 9f). In contrast with this successful observation, we also notified that magnified views of the skeletonized models show problems, such as, artificial excess connections of rhizoids forming of a loop-like structure (Fig. 9c) and artificial interruptions of a rhizoid (Fig. 9d).

**Fig. 9.**
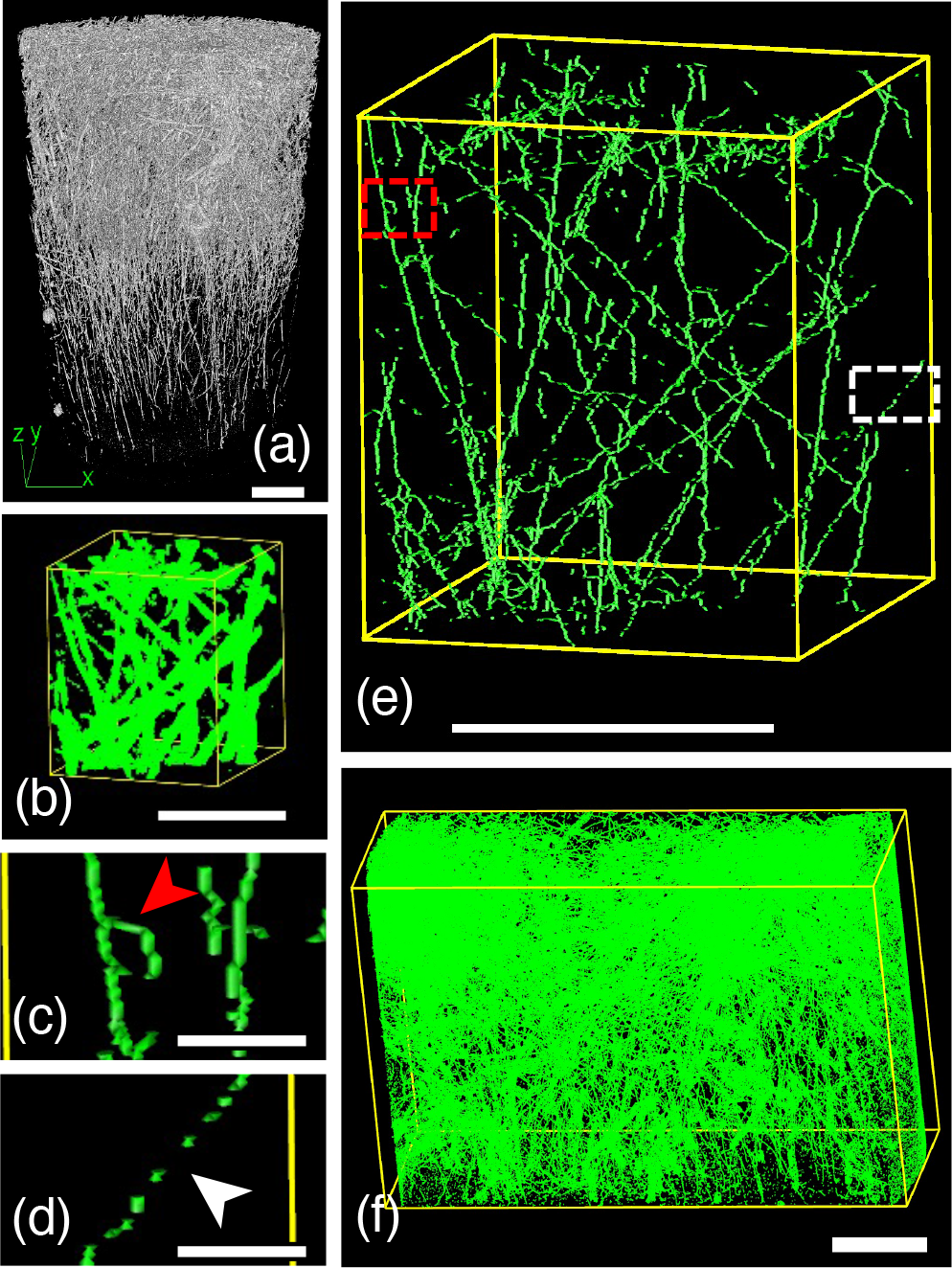
Three-dimensional models of the rhizoid system of *P. patens* embedded in paraffin. Isosurface models of the rhizoids (a, entire view; b, magnified view). Skeletonized individual rhizoid models (e). Magnified views of the skeletonized rhizoid models showing artificial formation of a loop-like structure (c) and artificial interruptions (d). A lower magnification view of skeletonized rhizoid models showing a part near the base of the rhizoid system (f). Models were visualized using 3D viewer of ImageJ software (a) or Imod software (b-f). Specimen example: Ground 1 × g_27 (a), Ground 1 × *g*_82 (b-f). Scale bars = 1000 μm (a, f), 50 μm (b, c, d), 500 μm (e).

## Discussion

To capture 3D morphology of the rhizoid system upto centimeter-order depth, specimens were processed for paraffin embedment as the first choice considering its versatility as well as processability: ease of trimming of centimeter-size block. On top of that, we have successfully found that automatic segmentation of rhizoid area based on machine learning (TWS 3D) method enabled to label the rhizoids, which appeared in different three form types in tomograms. In contrast, unfortunately, we had not noticed possible artifacts of paraffin embedment during preparation of Space Moss experiment performing optical microscopy of deparaffinized sections. Therefore, the rhizoid system specimens of Space Moss experiment prepared for X-ray micro-CT were embedded in paraffin and we have to use them for further analyses due to little opportunity for the next space experiment. It is also necessary to solve following problems, including the artifacts caused during paraffin embedment, for the future experiments.

### Technical problems regarding embedment of the rhizoid system

As far as the rhizoids become flat and/or void spaces are generated, quantitative analysis of the volume of the rhizoids is not possible because its volume would be underestimated when the rhizoids are flat and overestimated when they are replaced to void spaces.

The artifacts possibly due to paraffin embedment of samples, i.e., flattening of the rhizoids and generation of a void space at the location of the rhizoid, were noticed during 3D observations by X-ray micro-CT while these were not during preparatory observation of paraffin section before performing X-ray micro-CT. Our unawareness of this problem was possibly because the rhizoid tended to be flat after deparaffinization of the paraffin sections because the rhizoids frequently ran obliquely in the section. The reason why the rhizoid cell walls appeared thicker in the paraffin sections is also the same.

Flattening of the rhizoids occurred during paraffin embedment is possibly due to difficulty of infiltration of paraffin into the rhizoid cells. We have tested stirring and increasing of substitution steps (%) of paraffin during paraffin infiltration though it is still preliminary. However, these trials were not successful in avoiding of the artifact of rhizoid flattening. Using vacuum oven during paraffin infiltration might help to avoid the artifact of void space formation. On the other hand, embedment of the whole rhizoid systems in LR White resin is highly worth testing to avoid these artifacts.

### Issues regarding automatic segmentation

TWS 3D provides supervised machine-learning algorithms and thus prediction results may vary depending on the number of epochs (frequency of learning), accuracy for data annotation, and differences in preprocessing of the images. Optimizing these factors will increase accuracy of prediction, which remains to be an issue in the future.

The artifacts that became apparent after skeletonization are the artificial excess connection of the rhizoid (Fig. 9c) and the artificial interruption of the rhizoid (Fig. 9d). The former is considered to be due to excessive dilation and the latter is to excessive erosion for of the rhizoid area when Closing (3D) plugin software was applied. These problems may have particularly occurred where densities of the rhizoids are high at, such as, near the base of the gametophore. These are conflicting problems but need to be solved. Providing that current problems regarding artificial excess connection and artificial interruption of the rhizoids are minimized before skeletonization, the length of the rhizoids can be quantitatively analyzed more precisely.

Nevertheless, machine learning-based skeletonized 3D model revealed quite dense distribution of rhizoids near the base (Fig. 9f), and distribution density of rhizoids appears to decrease with distance from the base of the rhizoid system (Fig. 9a). While an overview of the entire architecture shows positive orthogravitropic response (i.e., growth oriented along the gravity vector) of the rhizoid system as a whole (Fig. 9a), it is also noticed that many rhizoids actually show plagiogravitropic responses (i.e., growth oriented obliquely relative to the gravity vector) in a magnified view (Fig. 9e). This is somehow similar to the root system architecture in vascular plants where a combination of orthogravitropism and plagiogravitropism of individual roots results in their wide distribution as a whole.

### Conclusions

The rhizoids of *P. patens* having the diameter of 21.3 μm on the average were visualized by refraction-contrast X-ray micro-CT using coherent X-ray optics available at the beamline of the synchrotron radiation facility SPring-8. Filamentous structures which appeared in different three form types in tomograms reconstructed from specimens embedded in paraffin were confirmed to be the rhizoids by optical and electron microscopy. Comprehensive automatic segmentation of the rhizoids, which appeared in different three form types in tomograms, was tested by image processing of contour extraction using Canny edge detector or that based on machine learning. Performance of the automatic segmentation methods was evaluated by comparing with the ground truth images using measures such as F1 score and IoU, revealing that the automatic segmentation method based on machine learning (TWS 3D) is more effective than that using Canny edge detector. Isosurface and skeletonized models of the rhizoid system automatically segmented using machine learning were visualized. This is the first time to successfully visualize the moss rhizoid system in 3D.

## Acknowledgements

The synchrotron radiation experiments were performed at the BL20B2 of SPring-8, with the approval of the JASRI (Proposal Nos. 2020A1264 and 2021B1316).

---

**Supplementary Fig. 1.** Movie made from consecutive CT tomographic slices in z direction. Yellow rectangular shows a filamentous structure which changes its form Type A, C, A, to B consecutively. Specimen example: Ground 10 × *g*_2. Scale bar = 50 μm.

**Supplementary Fig. 2.** Evaluation of the automatic segmentation methods. Rhizoid (white) and background (black) areas obtained by manual segmentation (ground truth) (a) and automatic segmentation (b) were compared and categorized into the areas of four groups: true positive (orange, TP), false positive (blue, FP), false negative (green, FN), and true negative (gray, TN) (c). A confusion matrix was constructed by counting the number of pixels in each categorized area (d). (e) Formulas calculating accuracy, F1 score, and Intersection over Union (IoU).

