## Supplementary Fig. 1 for "Three-dimensional visualization of moss rhizoid system by refraction-contrast X-ray micro-computed tomography"

### Slide 1
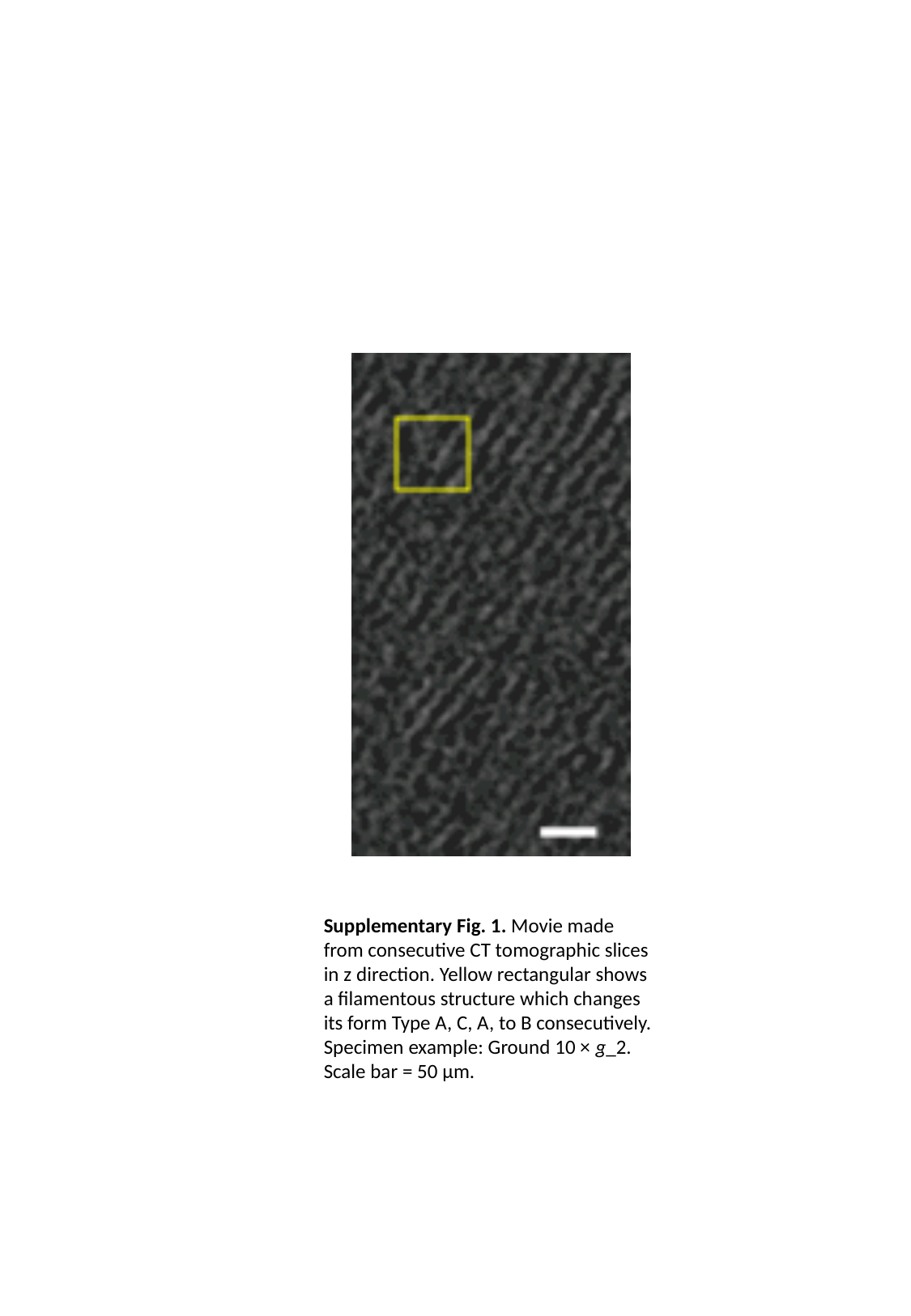

Supplementary Fig. 1. Movie made from consecutive CT tomographic slices in z direction. Yellow rectangular shows a filamentous structure which changes its form Type A, C, A, to B consecutively. Specimen example: Ground 10 × g_2. Scale bar = 50 µm.
