## Supplementary Fig. 2 for "Three-dimensional visualization of moss rhizoid system by refraction-contrast X-ray micro-computed tomography"

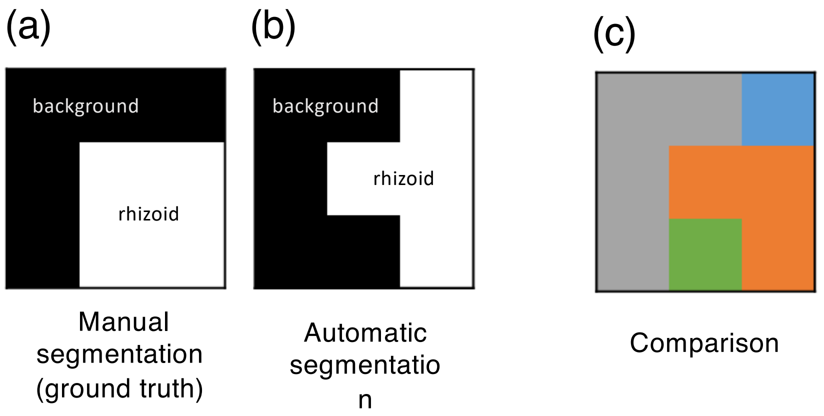


(d)

| Confusion matrix classification | | Predicted image | |
|---|---|---|---|
| | | Rhizoid | Back-ground |
| Actual (Ground Truth) image | Rhizoid | TP | FN |
| | Back-ground | FP | TN |

(e)

$$Accuracy = \frac{TP+TN}{TP+FP+FN+TN} \quad \cdots (1)$$

$$F1\ score = 2 * \frac{(Precision * Recall)}{Precision + Recall} \quad \cdots (2)$$

$$Precision = \frac{TP}{TP+FP}, \quad Recall = \frac{TP}{TP+FN} \quad \cdots (3)$$

$$IoU = \frac{TP}{TP+FP+FN} \quad \cdots (4)$$

**Supplementary Fig. 2**. Evaluation of the automatic segmentation methods**.** Rhizoid (white) and background (black) areas obtained by manual segmentation (ground truth) (a) and automatic segmentation (b) were compared and categorized into the areas of four groups: true positive (orange, TP), false positive (blue, FP), false negative (green, FN), and true negative (gray, TN) (c). A confusion matrix was constructed by counting the number of pixels in each categorized area (d). (e) Formulas calculating accuracy, F1 score, and Intersection over Union (IoU).
